# Neuroprotective strategies for NMDAR-mediated excitotoxicity in Huntington’s Disease

**DOI:** 10.1101/076885

**Authors:** KD Girling, YT Wang

## Abstract

**BACKGROUND:** Huntington’s Disease (HD) is an autosomal dominant neurodegenerative disease causing severe neurodegeneration of the striatum as well as marked cognitive and motor disabilities. Excitotoxicity, caused by overstimulation of NMDA receptors (NMDARs) has been shown to have a key role in the neuropathogenesis of HD, suggesting that targeting NMDAR-dependent signaling may be an effective clinical approach for HD. However, broad NMDAR antagonists are generally poor therapeutics in clinical practice. It has been suggested that GluN2A-containing, synaptically located NMDARs activate cell survival signaling pathways, while GluN2B-containing, primarily extrasynaptic NMDARs trigger cell death signaling. A better approach to development of effective therapeutics for HD may be to target, specifically, the cell-death specific pathways associated with extrasynaptic GluN2B NMDAR activation, while maintaining or potentiating the cell-survival activity of GluN2A-NMDARs.

**OBJECTIVE:** This review outlines the role of NMDAR-mediated excitotoxicity in HD and overviews current efforts to develop better therapeutics for HD where NMDAR excitotoxicity is the target.

**METHODS:** A systematic review process was conducted using the PubMed search engine focusing on research conducted in the past 5-10 years. 250 articles were consulted for the review, with key search terms including “Huntington’s Disease”, “excitotoxicity”, “NMDAR” and “therapeutics”.

**RESULTS:** A wide range of NMDAR excitotoxicity-based targets for HD were identified and reviewed, including targeting NMDARs directly by blocking GluN2B, extrasynaptic NMDARs and/or potentiating GluN2A, synaptic NMDARs, targeting glutamate release or uptake, or targeting specific downstream cell-death signaling of NMDARs.

**CONCLUSION:** The current review identifies NMDAR-mediated excitotoxicity as a key player in HD pathogenesis and points to various excitotoxicity-focused targets as potential future preventative therapeutics for HD.

## Huntington’s Disease

Huntington’s Disease (HD) is an autosomal-dominant, inherited neurodegenerative disease caused by a CAG expansion (>36 copies) in the huntingtin gene, leading to expression of a huntingtin (Htt) protein with an expanded polyglutamine tract near the N-terminus (Huntington’s Disease Collaborative Research Group, 1992). Early symptoms of HD include cognitive disruptions and alterations in mood, later developing to dance-like motor dysfunction (chorea), substantial neurodegeneration, dementia and eventually dyskinesia (Tobin 2003). Typically, symptomatic onset of HD occurs between the ages of 35-55, with death occurring 10-15 years after onset (Hayden 1981). Age of HD onset is linked to CAG repeat length and with longer CAG repeats (>60), juvenile HD may occur, with bradykinesia, rigidity, dystonia, cognitive changes and epileptic seizures, often without chorea (Huntington’s Disease Collaborative Research Group, 1992). The protein huntingtin (Htt) is ubiquitously expressed throughout the body and tissues, however pathology of HD is primarily brain specific, with most profound degeneration occurring in the striatum (Vonsattel et al. 1985). GABA-ergic medium spiny neurons (MSNs), constituting about 90-95% of striatal neurons, are particularly vulnerable in HD, whereas aspiny cholinergic interneurons are relatively spared (Ferrante et al. 1987). Although neurodegeneration is most prominent in striatal MSNs, non-autonomous cell death and dysfunction can also be detected in the cerebral cortex, globus pallidus, substantia nigra, white matter and hippocampus, and there is evidence of some damage to peripheral tissues and organs, including skeletal muscle (Spargo et al. 1993; Heinsen et al. 1996; Kassubek et al. 2004, (Abildtrup and Shattock 2013; Rosas et al. 2003; Ehrlich 2012). Despite a known genetic cause, treatments for HD are, at present, primarily palliative and there remains a lack of effective preventative HD therapies. Studies investigating preventative medicines for HD have targeted a wide variety of molecular mechanisms associated with cell death and dysfunction, including mitochondrial dysfunction (Tabrizi et al. 2000, Lim et al. 2008), caspase activation and cleavage (Wong et al. 2015; Graham et al. 2011; Hermel et al. 2004; Wellington et al. 2002; Carroll et al. 2011; Graham et al. 2006; Uribe et al. 2012;Graham et al. 2010) and BDNF dysregulation (Strand et al. 2007; Zuccato et al. 2010).

## The glutamate excitotoxicity hypothesis of HD

One major hypothesis as to the specific vulnerability of the striatum in HD is called the **excitotoxicity hypothesis** (Fan and Raymond 2007). Striatal neurons receive glutamatergic input from several sources, importantly from the cortex and thalamus which stimulated glutamate receptors on striatal MSNs. Several decades of research have demonstrated that excessive glutamatergic stimulation of these receptors via impaired uptake, enhanced glutamate release, enhanced sensitivity of the receptors or impaired downstream signaling of glutamate receptors may contribute in important ways to striatal vulnerability in HD (Schwarcz et al. 1977; DiFiglia 1990). In this review we will overview the role of NMDARs in glutamate excitotoxicity, investigate evidence for NMDAR-mediated excitotoxicity in HD pathogenesis and outline current neuroprotective strategies for HD based on the excitotoxicity hypothesis.

## NMDA Receptor physiology

Glutamate is the primary excitatory neurotransmitter in the central nervous system (Kandel et al. 1995), exerting its actions either by activating metabotropic glutamate receptors (mGluRs) which couple to G proteins, or via ionotropic glutamate receptors (iGluRs), which, upon ligand binding, allow passage of cations through a receptor pore (Watkins & Evans 1981; Dingledine et al. 1999). N-methyl D-aspartate (NMDA) receptors are the most highly studied of the iGluRs due to their importance in both normal neuron physiology and in disease pathology. To open, NMDARs require dual ligand binding of glutamate and the co-agonist glycine (Johnson & Ascher 1987) as well as removal of a Mg^2+^ block by membrane depolarization (Mayer et al. 1984). Activation of NMDARs causes influx of Ca^2+^ (MacDermott et al. 1986) which activates signal transduction cascades. The slow activation and deactivation kinetics of NMDARs govern the duration of the excitatory post-synaptic potential (Lester et al. 1990), giving NMDARs an important role in synaptic strength modulation. Native NMDARs are tetrameric complexes of two GluN1 (NR1) with two GluN2 (NR2) and/or GluN3 (NR3) subunits (Benveniste & Mayer 1991; Clements & Westbrook 1991). Different NMDAR subunit combinations change the ion properties and pharmacology of the receptor allowing for wide functional diversity of NMDARs (Monyer et al. 1992; Ishii et al. 1993; Flint et al. 1997; Chen et al. 1999). GluN2 subunits in particular are encoded by four genes (GluN2A-D) which determine differences in NMDAR channel properties, pharmacology and distribution (Dingledine et al. 1999; Cull-Candy & Leszkiewicz 2004). GluN2 subunits are both spatially and developmentally regulated (Monyer et al. 1992; Akazawa et al. 1994). In the forebrain of adults, the majority of NMDARs contain GluN2A and GluN2B subunits, with most NMDARs being biheteromeric GluN1-GluN2A or GluN1-GluN2B or triheteromeric GluN1-GluN2A-GluN2B receptors (Sheng et al. 1994; Li et al. 1998; Chapman et al. 2003). NMDA receptors are largely located at synaptic sites, but can also found extra- or perisynaptically. During development, up to 1/3 of NMDARs are extra-synaptically located, while synaptic NMDARs slowly increase in proportion as the brain matures. However a significant proportion of NMDARs remain extrasynaptic during adulthood (Tovar et al. 2013; Rosenmund et al. 1993; Cottrell et al. 2000; Petralia et al. 2010).

## NMDAR-mediated excitotoxicity: The role of glutamate

NMDARs have an important role in many neurological functions, including critical roles in synaptic plasticity, brain development and normal synaptic transmission (Aamodt & Constantine-Paton 1999; Bliss & Collingridge 1993). However, in many pathological conditions, overstimulation of NMDARs can trigger multiple neuronal death cascades, leading to apoptosis and necrosis (Berliocchi et al. 2005). This process is called **excitotoxicity** and has been implicated in a wide range of neuropathologies and neurodegenerative diseases including stroke (Lai et al. 2014. (Shiptoski 2012), Alzheimer’s Disease (Hynd et al. 2004; Koutsilieri and Riederer 2007), Parkinson’s Disease (Koutsilieri and Riederer 2007; Beal 1998) and neurotrauma (Marklund et al. 2004; Obrenovitch and Jutta 1997; Johnston 2005). Initial research on excitotoxicity arose from studies demonstrating that monosodium glutamate (MSG), an additive commonly found in Chinese food, was neurotoxic in the retina of the mouse (Lucas & Newhouse 1957). Subsequent studies demonstrated that this effect was not limited to the mouse retina, and similar effects were detected in both central and peripheral nervous system neurons in several other species (Burde et al. 1971; Freedman & Potts 1962; Olney & Sharpe 1969). Over several decades of research, it was discovered that the primary cause of neuronal death as a result of glutamate overstimulation was a result of excessive calcium influx (Choi 1995) primarily through NMDARs (Choi et al. 1988). Subsequently, a large number of studies have investigated the potential for NMDAR antagonists to protect against excitotoxic insults in various nervous system disorders, however the majority of NMDAR-antagonist studies fail to show efficacy in human clinical trials, due to multiple factors including side effects (Lipton 2004; Minnerup et al. 2012; Strokecenter.org 2016). The primary hypothesis for why NMDAR antagonists fail as therapeutics may be at least in part due to a paradoxical role of NMDARs, playing a pivotal role in both normal cellular function, including survival, and in neuronal death (Hardingham & Bading 2003). Blockade of all neuronal NMDARs using antagonists, though effective at preventing cell death pathways associated with NMDAR-mediated excitotoxicity also block the necessary, synaptic plasticity and cell survival pathways activated by NMDAR stimulation. These studies, and others, led to research into the underlying causes of the contradictory roles of NMDARs in hopes of developing better, more specific preventative therapies for nervous system disorders where NMDAR-mediated excitotoxicity has a role.

## A dichotomous role of NMDARs in cell survival and cell death

NMDARs are critical players in numerous functions related to cell survival and maintenance of neuronal homeostasis however NMDARs are also strongly involved in excitotoxic neuronal death. Although the underlying mechanisms of this dichotomous role of NMDARs is debated, there are many hypotheses as to the paradox of NMDARs. For one, the role of the NMDAR can vary depending on activity level. For example, when stimulated with low doses of NMDA, cultured granule cells show enhanced cell survival, however, with high dose stimulation with NMDA, the same neurons undergo cell death (Robert Balázs et al. 1988; Balázs et al. 1990; R. Balázs et al. 1988; Balázs et al. 1989; Didier et al. 1989; Yan et al. 1994). Similarly, cultured spinal cord neurons treated with low doses of NMDAR antagonist demonstrate enhanced cell survival, where the same neurons undergo cell death with high doses of the same antagonist (Brenneman, Forsythe, et al. 1990; Brenneman, Yu, et al. 1990). This, and other similar studies prompted an early hypothesis that there may be an optimal amount of intracellular calcium, and over or under stimulation of highly calcium permeable NMDARs leads to cellular death (Choi 1995; Koike et al. 1989; Franklin & Johnson 1992). In a similar vein, subsequent and emerging studies have demonstrated that both NMDAR location and receptor subunit composition may contribute to the differential role of the NMDAR in cell death and cell survival.

**Figure 1:**
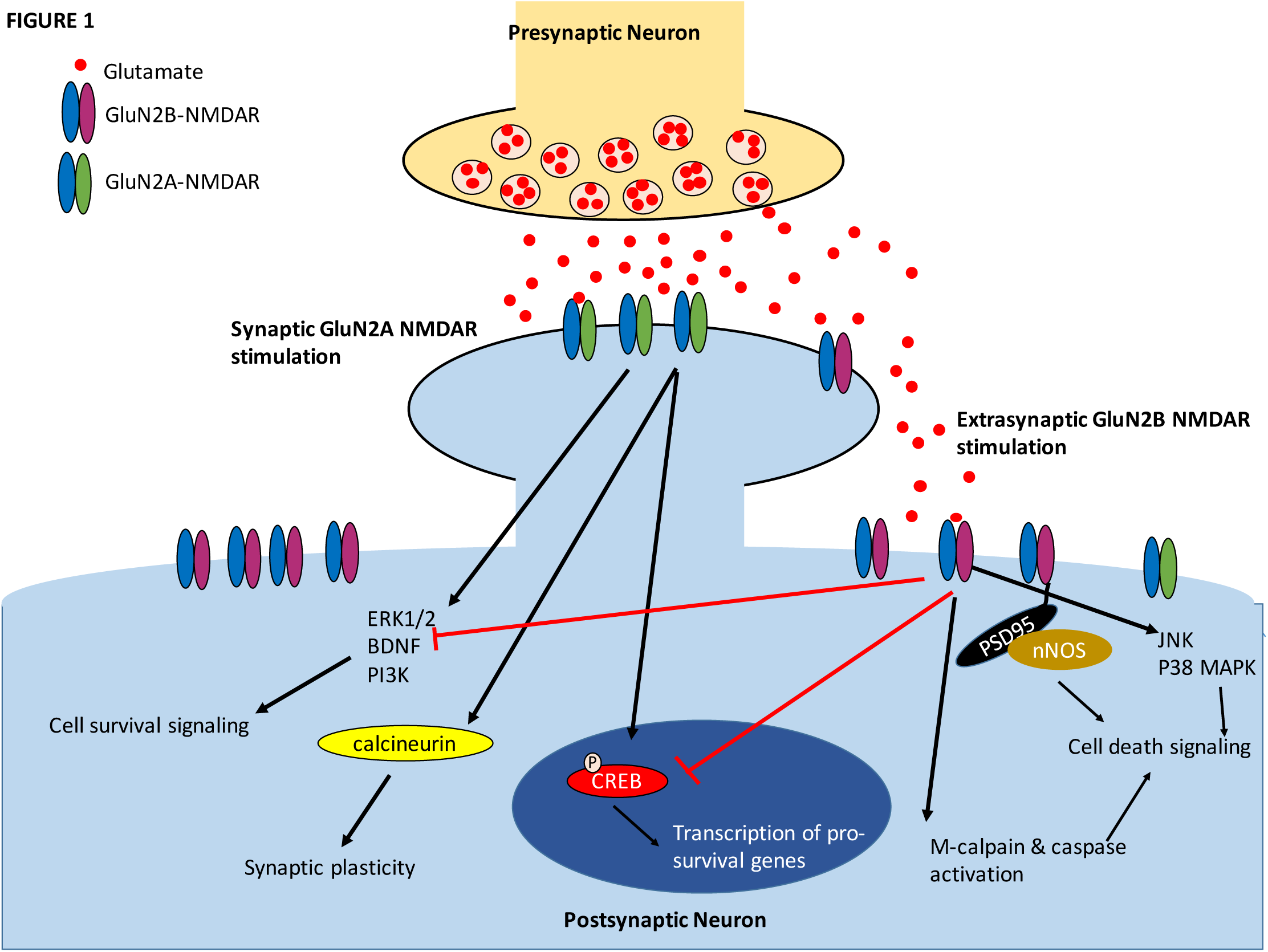
Dichotomous Role of NMDARs in cell survival and cell death. Activation of synaptically located, primarily GluN2A-containing NMDARs is associated with a wide range of cell survival and plasticity-promoting signaling pathways. Activation of these receptors leads to increased CREB phosphorylation in the nucleus, driving transcription of a range of cell survival-promoting genes. GluN2A, synaptic NMDAR activation also activates calcium-dependent calcineurin, shown to be involved in LTP and plasticity. Similarly, synaptic, GluN2A-NMDARs promote signaling of several cell survival molecular pathways including ERK1/2, PI3K and BDNF. Conversely, extrasynaptic, primarily GluN2B-NMDARs are related to cell death signaling. These receptors inhibit many of the synaptic, GluN2A pathways, including CREB shutoff, inhibition of ERK1/2, BDNF and P13K, and signals many molecular pathways involved in cell death, including JNK, and p38 MAPK. Extrasynaptic GluN2B NMDARs also activate cleavage molecules caspase and m-calpain, involved in protein cleavage, apoptosis and cell death.

## The subcellular location hypothesis for NMDAR-mediated excitotoxicity

One hypothesis for the dichotomous role of NMDARs in cell survival and cell death is that different populations of NMDARs may trigger different downstream signaling pathways upon activation. Many proteins with significant roles in molecular signaling and scaffolding are located exclusively at the synapse, such as PSD-95. Thus, it is likely that synaptically-located NMDARs, interacting with synapse-specific proteins, may behave differently than NMDARs located peri- or extrasynaptically. Many studies have demonstrated differences in downstream signaling pathways of synaptically located NMDARs compared to extrasynaptic NMDARs, prompting a theory of excitotoxicity called the “subcellular location” model These studies suggest that stimulation of synaptically-located NMDARs activates signaling pathways associated with cell survival and plasticity, whereas extrasynaptic NMDARs, trigger pathways associated with neuronal death (Lu et al. 2001; Hardingham & Bading 2003). This hypothesis was initially tested by selective enhancement of either synaptic or extrasynaptic NMDAR activity (Choi et al. 1988; Lu et al. 2001; Hardingham et al. 2002; Hardingham & Bading 2010). Synaptic NMDAR activity was enhanced pharmacologically, by blocking K^+^ channels using 4-aminopyridine, applying NMDAR co-agonist glycine which only enhances synaptic NMDARs that are activated by presynaptically released glutamate (Lu et al. 2001) or blocking GABAergic inhibition using bicuculline (Hardingham & Bading 2010), or by electrical stimulation, whereas extrasynaptic NMDARs were selectively stimulated using synaptic NMDAR blockade with MK-801, then bath application of NMDA(Hardingham & Bading 2010). Alternatively their contribution was attenuated by using the extrasynaptic-preferential NMDAR antagonist memantine (Xia et al. 2010; Parsons & Raymond 2014). Synaptic NMDARs stimulation causes a calcium-dependent upregulation of several pro-survival genes, including several anti-apoptotic factors, and suppression of several genes involved in cell death, leading to enhanced neuroprotection, reduced apoptotic ability and stimulating innate antioxidative properties of the cell (Hardingham et al. 2002; Hardingham & Bading 2010; Parsons & Raymond 2014). This synapse-specific NMDAR activity subsequently drives many cell-survival pathways, including extracellular signal-related kinase 1/2 activation, cAMP response elevated-binding protein (CREB) phosphorylation and enhanced expression of brain-derived neurotrophic factor (BDNF) (Hardingham et al. 2002; Hardingham & Bading 2010; Xu et al. 2009). On the contrary, specific stimulation of extrasynaptic NMDARs activates several molecular pathways that drive cell death, including CREB shut off, ERK1/2 inactivation and enhanced gene expression and activation of pro-apoptotic proteins such as Forkhead box protein O (Hardingham et al. 2002; Hardingham & Bading 2010; Xu et al. 2009). Similarly, calpains are differentially regulated by synaptic and extrasynaptic NMDARs. Synaptic NMDARs lead to activation of µ-calpain, whereas extrasynaptic NMDARs specifically activate m-calpain (Wang et al. 2013; Parsons & Raymond 2014),. m-calpain is preferentially involved in the cleavage of striatal-enriched tyrosine phosphatase (STEP), which subsequently activates p38 mitogen-activated protein kinase (p38MAPK) resulting in increased cell death (Hardingham et al. 2002; Hardingham & Bading 2010; Xu et al. 2009). In line with this data, stimulation of synaptic NMDA receptors is neuroprotective against neuronal insult caused by starvation or staurosporine (Hardingham et al. 2002; Papadia et al. 2005) whereas global NMDAR stimulation leads to neuronal death (Hardingham et al. 2002; Gouix et al. 2009; Zhang et al. 2007).

**Figure 2:**
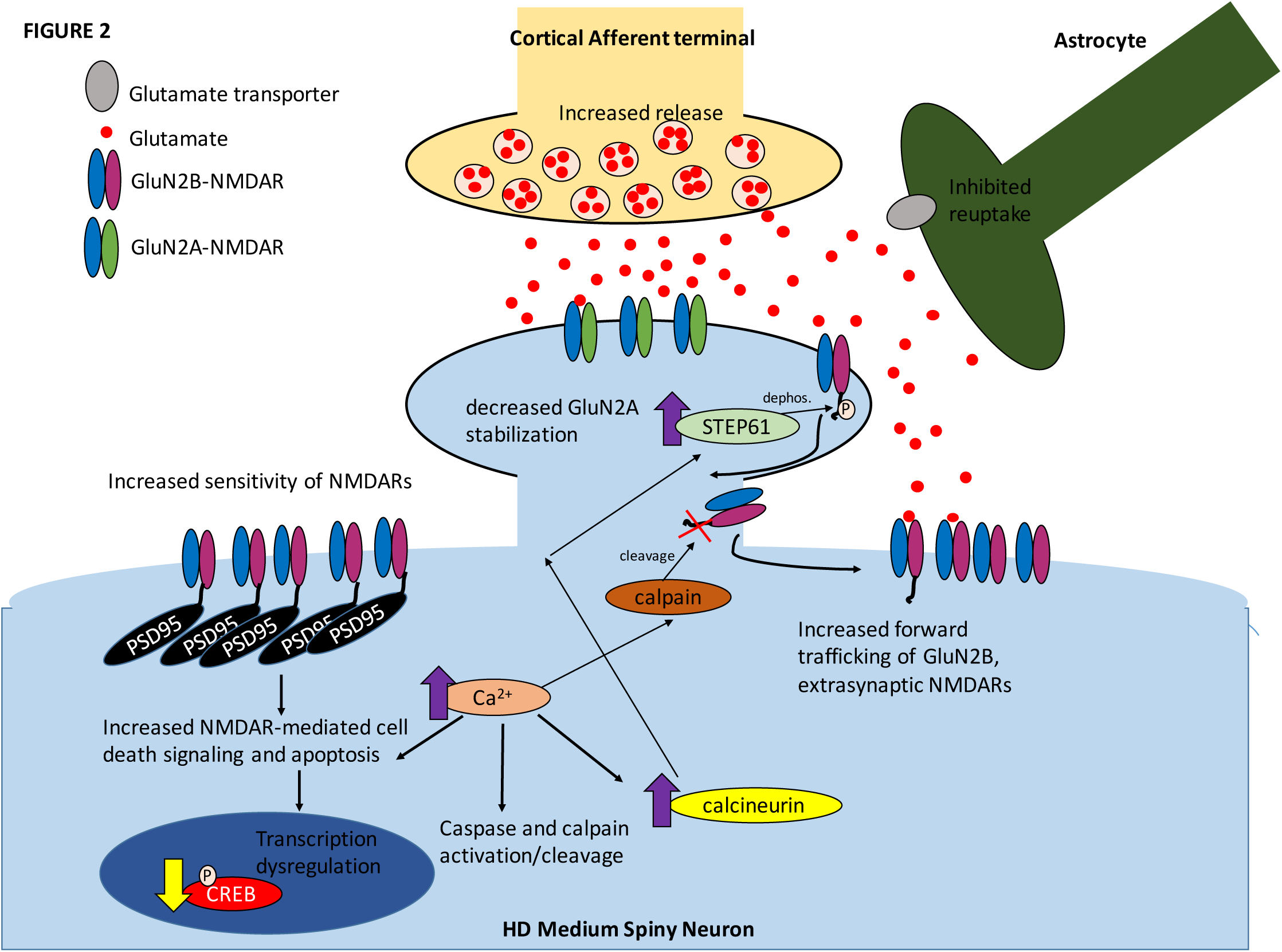
Schematic diagram of mHTT-dependent dysregulation in NMDAR expression and signaling in HD. Increased release from cortical (or thalamic) afferents stimulates NMDARs on medium spiny neurons in the striatum. Increased release and reduced reuptake from astrocytes leads to increased glutamate at the synapse, causing spillover and stimulating extrasynaptic, GluN2B-containing NMDARs, shown to be associated with cell death signaling. Increased NMDAR stimulation leads to increases in intracellular calcium, which stimulates increase in apoptotic signaling pathways, as well as activation of calcium-dependent molecules, such as caspases and calpains, which cleave mHTT into toxic fragments and stimulate cell death signaling. Calpain activation also leads to increased cleavage of GluN2B NMDARs, facilitating their movement from synaptic to extrasynaptic sites. Ca2+ also activates the calcium dependent molecule calcineurin, which subsequently activates STEP, which dephosphorylates and destabilizes synaptic NMDARs, reducing their expression. STEP also leads to the clathrin vesicle endocytosis of NMDARs, and their recycling to extasynaptic sites (not shown). mHTT influences PSD-95 interaction with NMDARs, interacting less strongly with GluN2A-NMDARs in the synapse, while stabilizing GluN2B-NMDARs extrasynaptically. Increased expression and stabilization of extrasynaptic NMDARs leads to increased sensitivity to NMDA, increased cell death and apoptotic signaling and dephosphorylation of CREB in the nucleus, leading to transcriptional dysregulation.

Synaptic and extrasynaptic NMDARs have also been shown to have opposite roles in regulation of synaptic plasticity (Lu et al. 2001). It is well known that NMDARs have an essential role in both long term potentiation (LTP) and long term depression (LTD), however recent data has demonstrated that increasing extracellular glutamate leads to impairments in LTP (Izumi et al. 2008; Katagiri et al. 2001; Li et al. 2011), an effect that was reversible using NMDAR antagonists ((Izumi et al. 2008; Katagiri et al. 2001; Li et al. 2011). Similarly pre-blockade of synaptic NMDARs before bath NMDA application leads to LTD formation (Liu et al. 2013). These, and other data, suggest that synaptic NMDARs may be preferentially involved in LTP formation, whereas extrasynaptic NMDARs are necessary to facilitate LTD. Taken together, a wide range of data suggests that subcellular location of NMDARs may have large impact on the differential effects of these receptors with synaptic NMDARs preferentially involved in neuron survival machinery, and extrasynaptic NMDARs facilitating neuronal death.

## The subunit hypothesis of NMDAR-mediated excitotoxicity

Another hypothesis for the paradoxical effects of NMDARs suggests that the physiological makeup of NMDARs contributes to the differential effects of the receptor. Most NMDARs are composed of two essential GluN1 subunit(Moriyoshi et al. 1991; Yamazaki et al. 1992) and two GluN2 subunits(Kutsuwada et al. 1992; Yamazaki et al. 1992; Mori & Mishina 1996). Variations in the GluN2 subunit leads to variations in receptor kinetics, properties and downstream signaling pathways due to difference in the carboxy-terminus of the receptor (Groc et al. 2006; Martel et al. 2009; Sanz-Clemente et al. 2013). Similar to the location hypothesis of NMDAR-mediated excitotoxicity, it has been proposed that subunit composition of NMDARs may underlie differential effects of NMDARs on cell survival and cell death. Using recently developed agonists and antagonists for GluN2A- and GluN2B-containing NMDARs, as well as genetic deletion of receptor subtypes, researchers have been able to investigate the particular role of each receptor subtype on neuron function, as well as the role in neuron survival vs death, and as a result, there has been support for a hypothesis that GluN2A-containing NMDARs have a primary role in cell survival signaling, whereas GluN2B-containing NMDARs are largely involved in cell death signaling (Liu et al. 2007; Zhou & Baudry 2006; DeRidder et al. 2006). Using specific antagonists for GluN2A or GluN2B-containing NMDARs, it has been shown that specific stimulation of GluN2B-containing NMDARs triggers excitotoxic neuronal death and apoptotic cascades, while stimulation of GluN2A-containing NMDARs is neuroprotective both in vitro against NMDA and non NMDAR-mediated neuronal death, as well as *in vivo* (Liu et al. 2007; Terasaki et al. 2010; Lai & Wang 2010) Similarly, specific stimulation of GluN2A-NMDARs is associated with activation of several downstream signaling pathways associated with cell survival such as CREB (Liu et al. 2007), P13K and kinase-D-interacting substrate of 220 kDa kinase-D-interacting substrate of 220 kDa (Kidins220) (López-Menéndez et al. 2009), whereas GluN2B-NMDAR stimulation activates numerous cell death specific pathways (Lai & Wang 2010; Martin & Wang 2010). Further, it has been shown that the subtype-specific differences in function of NMDARs are conferred by differences in the C-terminus of the NMDAR (Foster et al. 2010; Sprengel et al. 1998). To further test the differential role of NMDAR subtype on cell death and survival, Martel et al performed an experiment in which the C terminus of GluN2A was swapped with the C-terminus of GluN2B. In this experiment, the C-terminus of GluN2B enhanced neurotoxicity of the NMDAR when it replaced the GluN2A C-terminus (Martel et al. 2012), providing further evidence that GluN2B NMDARs are specifically involved in cell death, and GluN2A NMDARs enhance cell survival.

A controversial hypothesis of subunit-specific localization of NMDARs exists that suggests GluN2A-containing NMDARs are primarily located at synaptic sites in the forebrain, whereas GluN2B-containing NMDARs are mostly localized extra- and peri-synaptically (extrasynaptic sites) (Groc et al. 2006; Martel et al. 2009). This theory works in concert with location-based theories of NMDAR-mediated excitotoxicity, suggesting that synaptic/GluN2A NMDARs lead to cell survival while extrasynaptic/GluN2B-NMDARs enhance cell death. Much evidence has supported this theory over the years (Groc et al. 2006; Sanz-Clemente et al. 2013; Martel et al. 2009; Tovar & Westbrook 1999), however, there are several studies that have shown contradictory findings, and it is known that GluN2B can be found at synaptic sites and GluN2A can also be found outside of the synapse (Groc et al. 2006; Sanz-Clemente et al. 2013; Martel et al. 2009; Tovar & Westbrook 1999; Petralia et al. 2010; Liu 2004; Harris & Pettit 2007; Thomas 2006). Similarly, recent studies have shown some support for scenarios in which GluN2A, as well as synaptically located NMDARs can be implicated in cell death signaling under specific conditions (Papouin et al. 2012; Wroge et al. 2012; Zhou et al. 2013). These findings and others make the generation of a unified hypothesis for location and subcellular hypotheses of NMDAR-mediated excitotoxicity difficult to confirm. Similarly, variables such as the lack of very precise inhibitors for GluN2A and GluN2B, developmental changes in NMDAR subtype expression, growing knowledge of the importance of heterotetrameric NMDARs containing both GluN2A and GluN2B (Tovar et al. 2013) and the fact that a disproportionate percentage of NMDARS in cultured neurons are extrasynaptically located (Xia et al. 2010; Gladding & Raymond 2011) lead to challenges in determining the true underlying cause for differences in NMDARs in controlling cell survival and cell death.

Based on a wide breadth of research on potential causes for dichotomy in NMDARs, it is likely that a combination of subtype, subcellular location, developmental stage and type of activation lead to the varieties in NMDAR role in cell survival and cell death.

## NMDAR-mediated excitotoxicity in Huntington’s Disease

It has been widely demonstrated that NMDAR-mediated excitotoxicity has a critical role in the pathogenesis of Huntington’s Disease (HD). Evidence in human HD patients helped drive this theory, with the discovery that postmortem HD brains demonstrate reductions in NMDAR binding sites in the striatum (Albin et al., 1990; Young et al., 1988). Subsequently, brains of presymptomatic HD patients also showed increased expression in NMDARs in striatal MSNs, and it was found that these neurons were most vulnerable to death early on in disease progression (Graveland et al., 1985). Early studies using glutamate agonists quinolinic acid (QA) or kainic acid (KA) injected into the striatum of rats and later, to primates, led to neuropathological and behavioral changes similar to those seen in HD (Coyle and Robert 1976; Beal et al. 1991; Sanberg et al. 1989; Hantraye et al. 1990; Burns et al. 1995; Ferrante et al. 1993; McGeer and McGeer 1976; Schwarcz and Coyle 1977; Sanberg et al. 1978). Striatum neurons from pre-symptomatic transgenic HD mice show enhanced sensitivity to NMDA (Levine et al. 1999; Zeron et al. 2002), larger NMDAR currents (Zeron et al. 2002; Cepeda et al. 2001; Shehadeh et al. 2006), enhanced apoptotic death after NMDA stimulation (Shehadeh et al. 2006; Fan et al. 2007), increased surface expression of NMDARs (Shehadeh et al. 2006; Fan et al. 2007; Milnerwood et al. 2010), increased calcium responses during NMDA stimulation (Tang et al. 2005), as well as larger striatal lesions after intrastriatal QA (Graham et al. 2009; Hansson et al. 2001). This enhanced NMDAR excitotoxicity in HD appears to be mediated by extrasynaptic, GluN2B-containing NMDARs. Enhanced expression of extrasynaptic GluN2B-containing NMDARs are seen in HD transgenic mice (Milnerwood et al. 2010; Lipton 2004a). and a low dose memantine, blocking extrasynaptic NMDARS, is neuroprotective against neurodegeneration, synaptic dysfunction and behavioral dysfunction in mouse models of HD both *in vitro* and *in vivo* (Milnerwood et al. 2010; Lipton 2004a; Dau et al. 2014). Similarly, elevated striatal NMDAR current and excitotoxicity in primary striatum neurons and brain slices can be reversed with the GluN2B-selective antagonist ifenprodil (Zeron et al. 2002; Milnerwood et al. 2010; Fan et al. 2007; Tang et al. 2005).

Enhanced GluN2B expression and extrasynaptic localization in HD are, at least in part, regulated by enhanced forward trafficking and stabilization of GluN2B-NMDARs (Fan et al. 2007). The HTT protein has a wide variety of binding partners (Kaltenbach et al. 2007) many of which are disrupted by the presence of mHTT in HD (Zuccato et al. 2010). mHTT disrupts normal binding of HTT to PSD-95 (Sun et al. 2001) and subsequently enhances PSD-95 binding to GluN2B, believed to lead to enhanced stabilization of GluN2B NMDARs in HD (Fan et al. 2009; Milnerwood et al. 2010). mHTT-mediated impairments in clathrin-mediated endocytosis (Harjes & Wanker 2003), and endosomal receptor recycling (X. Li et al. 2009) as well as impaired phosphorylation of both mHTT and NMDARs (Gladding & Raymond 2011, Jarabek 2003; Lan et al. 2001) may also facilitate enhanced forward trafficking of NMDARs in HD (Gladding & Raymond 2011).

An important consequence of enhanced NMDAR-mediated excitotoxicity in HD is the activation of calcium-dependent calpain, shown to cleave NMDARs and affect receptor trafficking and diffusion between the synapse and extrasynapse (Guttmann et al. 2001; Guttmann et al. 2002). In HD, calpain activity is increased, leading to enhanced GluN2B cleavage (Cowan et al. 2008; Gafni & Ellerby 2002). Similarly, Striatal-Enriched protein tyrosine Phosphatase (STEP) has been shown to reduce synaptic NMDAR localization by dephosphorylation of GluN2B, and STEP activation is increased in early stage HD due to early increases in calcineurin activity (Paul et al. 2002). It has been recently suggested in a study using transgenic HD mice that these changes work together to enhance NMDA excitotoxic vulnerability in HD, with increased STEP61 activity leading to reductions in synaptic NMDARs while calpain cleavage of GluN2B enhances expression of extrasynaptic NMDARs (Gladding et al. 2012)

Although the striatum is the most widely studied brain area in HD, NMDAR dysfunction may also have a role in cognitive deficits in HD. Cognitive and mood disturbances are present in early stages of HD in humans and animal models, often preceding motor symptoms (Murphy et al. 2000; Van Raamsdonk 2005; Klapstein et al. 2001). Hippocampal loss is common in HD (Spargo et al. 1993; Usdin et al. 1999) and NMDAR-dependent hippocampal synaptic plasticity and transmission in CA1 pyramidal neurons is altered in early and presymptomatic stages in HD mice, suggesting a potential role for NMDARs in altered hippocampal function in HD (Murphy et al. 2000; Milnerwood et al. 2006; Milnerwood & Raymond 2007; Klapstein et al. 2001). Hippocampal neurons from YAC HD mice show hyper-excitability, reversed with NMDAR antagonists as well as increased resting cytosolic Ca2+ (Hodgson et al. 1999). Similarly, late stages of HD are accompanied by dementia and memory loss (Hansson et al. 2001; MacDonald et al. 1993). These data suggest that NMDAR excitotoxicity may also have a role in neuronal death and dysfunction in the hippocampus in HD, contributing to cognitive decline and memory deficit in HD.

Enhanced sensitivity to NMDA in HD models is present from birth and likely is an early underlying mechanism in HD pathogenesis (Zeron et al. 2002; Cepeda et al. 2001; Zeron et al. 2002; Cepeda et al. 2001; Shehadeh et al. 2006). Conversely, in later, symptomatic disease stages, HD transgenic mice of several strains develop resistance to NMDA both *in vitro (Hansson et al. 2001)* and *in vivo (Graham et al. 2009)* which may represent compensatory mechanisms in response to elevated calcium (Ca^2+^), reduced spine density(Hansson et al. 2001; Sun et al. 2002) or other neurological changes.

## Development of novel HD therapeutics targeting NMDAR-mediated excitotoxicity

Considering the prevalent role of NMDAR-mediated excitotoxicity in the pathogenesis of HD, many groups have investigated and given the ineffectiveness of broad NMDAR antagonists as therapeutics for excitotoxic neuronal death (Ikonomidou & Turski 2002; Lai et al. 2014; Tymianksi 2014). A wide array of studies have investigated potential ways to target cell death, specific death after an excitotoxic event, primarily by targeting GluN2B-containing, extrasynaptic NMDARs and the associated downstream signaling, while preserving GluN2A, synaptic NMDAR activity and signaling pathways. The following section will outline some of the major effort

## Targeting NMDA receptors

### Blocking GluN2B, extrasynaptic NMDAR activity

Given the broad evidence supporting the role of GluN2B-containing, extrasynaptically located NMDARs in HD, NMDAR antagonism seems a natural goal in the development of therapeutics. Early studies investigating excitotoxic insults in stroke and neurotrauma attempted to block NMDARs using broad NMDAR antagonists. Unfortunately, these drugs led to undesirable side effects in clinical application, largely because they also block the essential, neuroprotective role of NMDARs in normal functions including synaptic plasticity and cell survival (Ikonomidou & Turski 2002; Lipton 2004a; Lai et al. 2014, Kremer et al. 1999). Thus, developing potent and specific antagonist drugs for GluN2B-containing, extrasynaptic NMDARs is an area of intense interest. Ifenprodil and other similar drugs are a class of NMDAR antagonists showing selective, non-competitive binding for GluN2B NMDARs (Williams 1993; Fischer et al. 1997; Huang 1996; Kew et al. 1996). Initial experiments demonstrated effectiveness of these GluN2B antagonists in NMDAR-mediated excitotoxic models of stroke both *in vivo* and *in vitro* (Wang & Shuaib 2005; Liu et al. 2007; O’Donnell et al. 2006; Graham et al. 1992; Chen et al. 2008; DeRidder et al. 2006; Gotti et al. 1988) suggesting that they may be more therapeutically relevant drugs for excitotoxic conditions. Recent studies have demonstrated the potential therapeutic benefit of GluN2B antagonism in HD, using a co-cultured system of MSNs and cortical neurons, a more physiologically relevant culture system for studying glutamatergic synapses in HD (Milnerwood et al. 2012; Kaufman et al. 2012). Enhanced whole cell and extrasynaptic NMDAR-currents, increased sensitivity to NMDA and increased cell-death specific signaling can be detected in YAC128 HD MSNs compared to wild type in this system (Milnerwood et al. 2012). However, blocking GluN2B activity in these cultures using ifenprodil protects against NMDAR insults and mHTT-mediated CREB shutoff, whereas blocking GluN2A NMDARs does not (Milnerwood et al. 2012; Zeron et al. 2002). Similarly, enhanced NMDAR currents seen in MSNs from YAC HD mice can be attenuated using ifenprodil (Zeron et al. 2002; Milnerwood et al. 2010), as well as toxic mHTT nuclear inclusions (Okamoto et al. 2009) suggesting that GluN2B NMDAR antagonism may help alleviated neuropathological changes in HD. Despite promising effects of GluN2B antagonists in HD and other NMDA excitotoxic disease models, there are still limitations, namely 1) potential inefficacy of GluN2B antagonism without additional potentiation of GluN2A and subsequent cell survival signaling; 2) lack of strong subunit specificity of current GluN2B antagonists; 3) restrictive therapeutic window for GluN2B antagonists alone after an excitotoxic event (Yuan et al. 2015). These may help explain these limiting results. In addition, the dose-limiting side effects of GluN2B antagonism in clinical applications of excitotoxicity (Yuan et al. 2015) and potential for negative outcomes by blocking GluN2B, which also has a role in synaptic plasticity, could contribute to lack of efficacy in improving neurological outcomes. Despite promising effects of ifenprodil in the YAC128 HD model, GluN2B-specific antagonists have shown varied effects. For example, subcutaneous injections of three different GluN2B antagonists, ifenprodil, RO25,6981 and CP101,606, failed to show benefits in an R6/2 HD model *in vivo* (Tallaksen-Greene et al. 2010) suggesting that effective therapeutics for HD may need to expand upon GluN2B antagonism. Subsequent studies have attempted to improve efficacy of GluN2B antagonists in *in vivo* models of NMDAR-mediated excitotoxic neuronal death. A recent study attempted to maximize GluN2B antagonist effectiveness in ischemia based on the idea that ischemia is associated with acidification of tissues (Katsura et al. 1992) (Yuan et al. 2015; Matsumoto et al. 1990). Subsequently this study developed pH sensitive GluN2B antagonist compounds using medicinal chemistry to help limit NMDAR antagonism to ischemic tissue, while reducing effect in healthy brain (Yuan et al. 2015). These compounds provided significant neuropathological and behavioral improvements with minimal side effects (Yuan et al. 2015) suggesting that limiting GluN2B NMDAR antagonism to areas of the brain undergoing excitotoxicity may help mitigate some of the negative consequences of NMDAR antagonism, and pave the way to effective therapies.

Concurrent with GluN2B antagonism, another potential antagonism-based therapy for excitotoxicity in HD is memantine, the only current clinically-approved NMDAR antagonist. Memantine is a non-competitive NMDAR antagonist with fast on-off kinetics that, at low doses, has shown to preferentially block tonically-activated, extrasynaptically located NMDARs but not phasically activated synaptic NMDARs (Xia et al. 2010). Memantine is currently approved as a prescription medication for Alzheimer’s Diseases patients, and has shown to delay onset of behavioral and cognitive symptoms in human patients (Areosa Sastre et al. 1996; Howard et al. 2012). Given the critical role of extrasynaptic NMDARs in HD pathogenesis (Parsons and Raymond 2014; Kaufman et al. 2012; Milnerwood et al. 2012), researchers have investigated the potential of memantine to act as a therapeutic strategy against excitotoxicity in HD. Given at a low dose, shown to preferentially block extrasynaptic NMDARs, memantine was able to abolish the early sensitivity to NMDA seen in YAC128 HD mouse striatum *in vivo* (Okamoto et al. 2009; Okamoto et al. 2009). Memantine treatment also lead to motor improvement and reduced striatal loss (Okamoto et al. 2009; Milnerwood et al. 2010). More recently, memantine treatment in YAC128 HD mice was shown to also rescue synaptic dysfunction and downstream cell death signaling in HD, normalizing enhanced extrasynaptic NMDAR expression, reducing calpain activation, reducing p38 MAPK activation and rescuing CREB shutoff (Dau et al. 2014). Synaptic NMDAR activity and signaling were unaffected by low-dose memantine (Dau et al. 2014). While broad NMDAR antagonism isn’t ideal for HD therapeutic development, these studies demonstrate that disease- or area-specific GluN2B- or extrasynaptic-specific NMDAR antagonism may provide neuroprotective benefit in HD.

### Enhancing GluN2A, synaptic NMDAR activity

Based on evidence demonstrating that synaptically-located, GluN2A-containing NMDARs are preferentially tied to several fundamental cell-survival pathways (Hardingham & Bading 2003; Hardingham et al. 2002; Hardingham & Bading 2010; Martel et al. 2012; Lai et al. 2014; Liu et al. 2007) and that blockade of GluN2A-containing NMDARs using the antagonist NVP-AAV077 significantly worsens apoptosis, cell death and behavioral outcomes *in vitro* or *in vivo* models of stroke (Liu et al. 2007) a better option may be to enhance the function of GluN2A-containing, synaptic NMDARs to improve cell survival and function after excitotoxic insults. It has been shown that brief bath application of suprasaturating doses of glycine leads to selective stimulation of synaptically located NMDARs (Man et al. 2003; Lu et al. 2001). As glycine is a co-agonist of NMDARs, bath glycine application enhances activation of NMDARs selectively located in the synapse that are stimulated by spontaneous presynaptic glutamate release (Man et al. 2003; Lu et al. 2001) and not extrasynaptic NMDARs, which are not active in an unstimulated state (Man et al. 2003; Lu et al. 2001). Glycine enhancement of synaptic GluN2A-containing NMDARs leads to enhancement of cell survival signaling and reduces apoptotic cell death in *in vitro* NMDAR-mediated excitotoxicity (Liu et al. 2007). Similarly, in an MCAO stroke model in rats, glycine given post-stroke significantly reduced infarct size, an effect that was enhanced when animals were co-treated with GluN2B antagonist Ro-25–698 (Liu et al. 2007). One limitation of GluN2A agonism as a potential target for excitotoxic death is that there is a lack of specific, effective GluN2A agonists available. Similarly, despite many studies supporting a survival role of synaptic, GluN2A NMDARs, some studies refute this and suggest synaptic or GluN2A-containing NMDARs can also mediate cell death (Wroge et al. 2012, Papouin et al. 2012). However, based on data in excitotoxic stroke models, it is possible that compounds to enhance synaptic, GluN2A NMDARs may show neuroprotective benefit against NMDAR excitotoxicity and may be a promising next step in the search for better HD therapeutics, especially if used in conjunction with extrasynaptic GluN2B antagonism.

## Targeting glutamate release and uptake in HD

Another way in which striatum neurons may be particularly prone to NMDAR mediated excitotoxicity in HD is through impaired glutamate release and uptake. Though not observed in all mouse models (Li et al. 2004), some transgenic mouse models, such as the R6/2 HD model, show increased spontaneous EPSCs in striatum MSNs in acute slices in pre-symptomatic animals (Cepeda et al. 2003), indicative of enhanced glutamate release. Similarly, in HD patients and transgenic HD mice, metabolites 3-hydroxykyneurine, enhancing oxidative stress or quinolinic acid, stimulating NMDARs, are both augmented (Guidetti et al. 2006; Guidetti et al. 2004), further suggesting upstream changes that could enhance excitotoxic vulnerability in MSN NMDARs. Along these lines, impaired glutamate transport may also play a key role in HD excitotoxicity. Normally, glutamate transporters play an important role in preventing glutamate buildup at the synapse, reuptaking excess to prevent excitotoxicity. However, in HD, some studies have suggested that glutamate reuptake is impaired. In R6/2 HD mice, very early deficits in mRNA expression of the important glial transporter GLT-1 can be detected in both cortex and striatum tissues (Liévens et al. 2001) as well as decreased protein expression ((Faideau et al. 2010; Liévens et al. 2001). Impaired basal glutamate uptake by GLT-1 in HD mouse models has also been further reported by using microdialysis (Miller et al. 2008), measurement of impaired glutamate uptake in synaptosomes (Huang et al. 2010; Liévens et al. 2001) and D-aspartate binding (Liévens et al. 2001). Human HD brains similarly show impaired glutamate uptake using a [^3^H]-glutamate uptake assay or measuring [^3^H]-aspartate binding(Cross et al. 1986; Hassel et al. 2007). Subsequent studies have investigated the potential of reversing impairments in GLT-1 in HD models. The antibiotic ceftriaxone, shown to elevate expression of GLT-1, was recently used in R6/2 HD mice, given for 5 days (Miller et al. 2008). The drug effectively enhanced GLT-1 levels, reversed impaired glutamate uptake in HD animals vs wild type and led to improvements in HD phenotype (Miller et al. 2008). However, recent data using a more physiological *in situ* model of glutamate uptake demonstrated no impairments in glutamate clearance following synaptic release in YAC128 or R6/2 mice (Parsons et al. 2016).

## Targeting cell-death specific signaling of NMDARs

Given the ineffectiveness of broad NMDAR antagonists in clinical application, a potential therapeutic strategy for NMDAR-mediated excitotoxic death focuses on downstream pathways activated by NMDAR stimulation. As discussed, synaptic, GluN2A-NMDAR stimulation is specifically associated with many cell-survival signaling pathways. These pathways are antagonized during NMDAR-excitotoxicity by extrasynaptic GluN2B-NMDAR stimulation, which trigger downstream signaling associated with cell death. By developing therapeutic targets that specifically shut down cell death signaling associated with GluN2B, extrasynaptic NMDARs, the cell-survival signaling of synaptic, GluN2A NMDARs may remain intact, preventing the negative side effects of NMDAR antagonism.

## PSD-95

One way in which NMDARs confer differential outcomes on cell death and cell survival is by coupling directly with different interacting proteins. By interacting with proteins located specifically at synaptic or extrasynaptic locations, NMDARs may be functionally linked to different downstream signaling pathways for cell survival or cell death. Similarly, GluN2A and GluN2B NMDARs have shown to interact preferentially with cell survival and cell death-specific signaling molecules, respectively, by differential direct coupling via the NMDAR C-terminus (Foster et al. 2010; Sprengel et al. 1998; Martel et al. 2012). These important differences have allowed scientists opportunity to develop therapeutic potential drugs for NMDAR excitotoxic conditions that aim to disrupt direct or indirect NMDAR interaction with cell-death specific molecules, such as neuronal nitric oxide synthase (nNOS) (Aarts et al. 2002; Zhou et al. 2010; Sattler and Tymianski 2000), death-associated protein kinase 1 (DAPK1) (Tu et al. 2010; Fan et al. 2014) and PTEN (Zhang et al. 2013). A particularly important target of NMDAR-mediated interaction protein studies is PSD-95, a membrane-associated guanylate cyclase (MAGUK) that is found concentrated at the postsynaptic density of glutamatergic synapses, with essential roles in synapse stabilization and plasticity (El-Husseini et al. 2000). PSD-95 binds directly to GluN2B via PDZ domains (Kornau et al. 1995; Brenman et al. 1996), which has shown to facilitate excitotoxic death signaling, by linking GluN2B-NMDARs with nNOS (Aarts et al. 2002; Sattler and Tymianski 2000). mHTT has shown to exacerbate GluN2B-NMDAR interaction, with enhanced GluN2B-PSD-95 binding detected in striatal tissue from YAC transgenic HD mice at times when enhanced NMDA sensitivity is present (Fan et al. 2009), whereas this effect is gone later, when NMDA resistance occurs (Jarabek 2003). Uncoupling of PSD-95 with GluN2B using a small interfering peptide NR2B-9c (Aarts et al. 2002) reduced striatal sensitivity to NMDA to levels observed in WT MSNs (Fan et al. 2009). However, this mHTT-PSD-95-NMDAR mechanism in HD is thought to be independent of nNOS, and rather dependent on enhanced activation of p38 MAPK cell death signaling (Fan et al. 2012) as enhanced p38 MAPK can also be rescued with GluN2B-9c in HD transgenic models (Fan et al. 2012).

Another important role of PSD-95-GluN2B interaction in HD is through regulation of NMDAR trafficking and localization. PSD-95 interaction with NMDARs via the C-terminal tail has previously been shown to regulate NMDAR stabilization at synapses (Roche et al. 2001; Lin et al. 2004; Prybylowski et al. 2005). In the striatum of YAC transgenic models of HD altered NMDAR trafficking includes enhanced surface expression of NMDARs (Fan et al. 2009) as well as increased expression of PSD-95 at extrasynaptic sites (Milnerwood et al. 2010), shown to sensitize HD mice to NMDA excitotoxic cell death. Wild type huntingtin itself has also been shown to interact strongly with PSD-95 directly, linking it to NMDARs (Shirasaki et al. 2012); however, in HD, expanded polyglutamine weakens the Htt-PSD-95 interaction (Sun et al. 2001), leading to enhanced sensitivity of existing NMDARs to NMDA (Sun et al. 2001). Similarly, impaired mHTT-PSD-95 interaction increases extrasynaptic expression of PSD-95, a normally synapse-specific protein (Fan et al. 2012). Taken together, PSD-95’s strong interaction with GluN2B, strengthened by the presence of mHTT is thought to facilitate extrasynaptic expression of GluN2B-containing NMDARs by stabilizing them via PSD-95 (Fan et al. 2012; Parsons and Raymond 2014; Parsons et al. 2014). The NR2B-9c peptide further shows benefit to HD models by partially correcting mistaken trafficking of NMDARs, mildly reducing the enhanced surface expression of NMDARs in HD transgenic MSNs (Fan et al. 2009).

Several studies have investigated other potential ways to correct impaired PSD-95 and NMDAR localization in HD. Mislocalization of PSD-95 and extrasynaptic NMDARs in HD has been shown to be linked to post-translational modifications of NMDARs such as caspases, calpains, phosphorylation and palmitoylation (Gladding and Raymond 2011; Parsons and Raymond 2014). HD is linked with impairments in palmitoylation (Young et al. 2012; Sanders and Hayden 2015). Wild type HTT, along with its strong interaction with PSD, also interacts with the palmitoylacytransferase DHHC17, otherwise known as Huntingtin Interacting Protein 14 (HIP14) (Huang et al. 2011; Singaraja et al. 201; Sutton et al. 2013). In HD, mHTT-HIP14 interaction is impaired (Huang et al. 2011; Singaraja et al. 2011), and genetic knockout mice for HIP14 or the HIP14-like protein (HIP14L) share similar phenotypical features with HD (Singaraja et al. 2002). Palmitoylation of NMDARs can impact their surface expression and trafficking (Hayashi et al. 2009; Mattison et al. 2012), and similarly, palmitoylation of PSD-95 is important for synaptic targeting of the GluN2B-PSD-95 complex (Craven et al. 1999; Parsons and Raymond 2014). Along with HD-like phenotype, Hip14−/− mice also show reduced palmitoylation of PSD-95 (Singaraja et al. 2002). Current studies are investigating the role of impaired palmitoylation in HD on PSD-95 and NMDAR expression and localization.

## Calpains and STEP

Calcium influx as a result of NMDAR stimulation activates a wide variety of downstream molecular mechanisms. Calpains, a class of calcium-dependent cysteine proteases have been shown to have a role in NMDAR-mediated HD pathogenesis. Excitotoxic NMDAR activation of calpains (Siman & Carl Noszek 1988) has shown to have a key role in proteolysis, impaired synaptic function and neuronal damage after an NMDAR-mediated excitotoxic event (Arai et al. 1991; Lee et al. 1991; Arlinghaus et al. 1991; Lai et al. 2014; Rami & Krieglstein 1993). NMDAR-mediated calpain activity can also cause further calcium overload after NMDAR calcium entry, due to calpain-mediated cleavage of the sodium-calcium exchanger (NCX) (Bano et al. 2005), creating enhanced cell vulnerability to calcium-mediated cell death pathways (Lai et al. 2014). Calpain activation after NMDAR excitotoxicity appears to be GluN2B-specific, triggering cell death signaling such as the 35-kDa regulatory activator (p35) of cdk5 (Patrick et al. 1999; Lee et al. 2000), cleaving it into the toxic p25 fragment (Lee et al. 2000), and can be blocked using antagonists for GluN2B but not GluN2A (DeRidder et al. 2006;Gascón et al. 2008). Calpains are also differentially regulated by synaptic and extrasynaptic NMDARs, with synaptic NMDARs activating µ-calpain while extrasynaptic NMDARs activate m-calpain (Wang et al. 2013). m-calpain in particular cleaves striatal enriched tyrosine phosphatase (STEP), which subsequently activates p38-activated mitogen activated kinase (p38MAPK) (Wang et al. 2013; Xu et al. 2009), further contributing to differential NMDAR effects (Parsons and Raymond 2014).

In mouse models of HD, early, pre-symptomatic enhancement of calpain activation can be detected, along with a subsequent increase in the synaptic-specific STEP61 (Cowan et al. 2008; Gafni and Ellerby 2002; Gladding et al. 2012; Dau et al. 2014). Calpain and STEP61 activation significantly contributes to enhanced NMDA sensitivity in HD both by dephosphorylating the GluN2B NMDAR subunit, leading to enhanced expression of extrasynaptic and not synaptic NMDARs (Gladding et al. 2012). STEP61 also contributes to neuronal death in HD by dephosphorylation and deactivation of the survival-specific proteins ERK1/2 (Gladding et al. 2014) as well as STEP33 (a cleavage product of STEP61)-mediated activation of p38 MAPK, leading to p38 activation and subsequent cell death (Gladding et al. 2014). Activated p38MAPK can be detected in the striatum of transgenic HD mice both pre-symptomatically during enhanced sensitivity to NMDA, and later, NMDA resistant stages (Fan et al. 2012; Saavedra et al. 2011). Inhibition of calpain has shown to reverse the enhanced extrasynaptic NMDAR expression seen in the YAC128 HD model (Gladding et al. 2012; Gladding and Raymond 2011; Parsons and Raymond 2014). Calpain effects appear to be particularly important at early stages of the disease, as enhanced calpain is only detected at early, pre-symptomatic stages (Dau et al. 2014). These data point to calpain inhibition as potential early target to prevent enhanced sensitivity to NMDA as well as prevent p38 activation in later disease stages (Gladding et al. 2012). However, calpains are ubiquitously expressed and necessary for many aspects of cell survival (Lai et al. 2014) thus they should be viewed with caution when developing therapeutics.

## Caspases

A major downstream pathway associated with NMDAR-mediated excitotoxic and subsequent apoptotic death is the activation of caspases. Caspases are cysteine-aspartic proteases, activated as a part of the apoptotic cascade that act to post-translationally modify proteins by cleaving at specific sites, which may act as a gain- or loss- of function modification(Pop and Salvesen 2009). Caspases play an important role in apoptosis, and initiator caspases, activated both stress- and cell-type dependently, can initiate proteolytic caspase cascades, leading to recruitment and activation of executioner caspases and apoptotic cell death (Pop and Salvesen 2009; Graham et al. 2011). Caspase activation and apoptosis have shown to play an important role in HD pathogenesis. TUNEL staining in the affected areas of HD brains (Dragunow et al. 1995), indicates that apoptotic cell death contributes to HD neuropathy (Fan and Raymond 2007). Similarly, activated caspase-8 and caspase-6 can be detected in the tissues of transgenic HD animal models, as well as in the brains of both early and late stage HD patients (Sánchez et al. 1999; Graham et al. 2006; Hodges et al. 2006; Graham et al. 2010; Hermel et al. 2004) and increased proform casp3 can be detected in later stage HD human brain tissue (Graham et al. 2010). The huntingtin protein contains several consensus sites for caspase cleavage (Wellington et al. 1998; Wellington et al. 2000) and is proteolytically cleaved by caspases in both HD and normal tissues. In HD, caspase cleavage of the mutated mHTT creates toxic fragments, shown to have a key role in HD pathogenesis both *in vitro* and *in vivo* (Graham et al. 2006; Graham et al. 2010; Wellington et al. 2002; Mangiarini et al. 1996). Caspase-mediated apoptosis in HD appears to be NMDA excitotoxicity-dependent. Cultured MSNs from transgenic YAC128 HD mice demonstrate enhanced casp6 mRNA (Graham et al. 2006) and cultured YAC42 and YAC72 HD striatum neurons have increased levels of active casp3 after NMDA stimulation (Zeron et al. 2002). Similar increases in casp3 are detected in human HD lymphoblasts when stimulated with mitochondrial stressors(Sawa et al. 1999). Using fluorescence substrate cleavage assays *in vitro*, it has been demonstrated that caspase activation effects in HD are mediated by the intrinsic apoptotic cascade, rather than the extrinsic death-receptor mediated pathway (Zeron et al. 2004).

Several studies have investigated the potential for caspase inhibition as a means of protecting against HD-induced neuropathologies. In cultured HD neurons, inhibition of casp3, casp6 or casp9 protects cells against NMDAR-mediated apoptotic death (Tang et al. 2005; Graham et al. 2006). Similarly, minocycline treatment, reducing casp1 and casp3 mRNA, slows HD progression in mouse HD models *in vivo* (Chen et al. 2000). However, casp3 inhibition alone doesn’t work in all HD models (Kim et al. 1999). Casp6, in particular, has been proposed as an initiator caspase in apoptotic HD cell death cascades (Graham et al. 2011). Levels of activated casp6 correlate positively with the size of the CAG repeat and inversely with age of HD onset (Graham et al. 2010). Transgenic HD mice engineered with a casp6 resistant mHTT (C6R mice) show neuroprotection against HD striatal atrophy and neurodegeneration (Graham et al. 2006). Similarly, C6R striatal neurons show neuroprotection against NMDA and staurosporine *in vitro* and C6R mice have reduced quinolinic acid-induced striatal lesions, improved behavioral outcomes and improvements in neurological HD changes (Graham et al. 2006; Pouladi et al. 2009; Warby et al. 2008; Graham et al. 2011; Graham et al. 2010; Milnerwood et al. 2010; Wellington et al. 2002). Similarly, chemical casp6 inhibition (Graham et al. 2010), dominant-negative caspase inhibition (Hermel et al. 2004) or genetic silencing of casp6 (Uribe et al. 2012;Wong et al. 2015) are neuroprotective and behaviorally beneficial in several models of HD. These data, as well as data demonstrating early activation of casp6 in pre-symptomatic and early stage HD brains, while casp3 isn’t active until later in disease progression (Graham et al. 2010) suggest that casp6 may be activated early, followed by casp3 activation, and may be a possible therapeutic target for HD (Graham et al. 2011).

## Conclusions

NMDAR-mediated excitotoxicity has been shown to have a critical role in the pathogenesis of Huntington’s Disease. Thus, the understanding of pathways involved in cell death after excitotoxicity is of utmost importance. Development of effective and clinically applicable drug targets for NMDAR-mediated excitotoxicity in HD have shown potential benefits in a wide range of animal models and may represent novel therapeutics for HD that are aimed at presymptomatic prevention of symptoms, rather than palliative treatments. Similarly, developing novel drugs for NMDA exitotoxicity meets a large goal in the medical community as they may have wide reaching applicability to not only HD, but also use in many neurodegenerative diseases of aging and brain injury.

## Compliance with Ethical Guidelines

1. Conflict of Interest Kimberly Girling and Yu Tian Wang declared that they have no conflict of interest.
2. Human and Animal Rights, and Informed Consent All institutional and national guidelines for the care and use of laboratory animals were followed.

This manuscript is a review article and does not involve a research protocol requiring approval by the relevant institutional review board or ethics committee.

